# *Vibrio cholerae* O1 El Tor A1552 encodes two functional ornithine lipid synthases and induces ornithine lipid formation under low phosphate and under low salinity growth conditions

**DOI:** 10.1101/2024.12.11.627999

**Authors:** Miguel Ángel Vences-Guzmán, Roberto Jhonatan Olea-Ozuna, Raquel Martínez-Méndez, Wendy Itzel Escobedo-Hinojosa, Marlene Castro-Santillán, Ziqiang Guan, David Zamorano-Sánchez, Christian Sohlenkamp

## Abstract

Ornithine lipids (OLs) are phosphorus-free membrane lipids that can be formed by a wide range of bacteria. The presence of OLs is frequently related to the resistance to abiotic stress conditions, and its synthesis is often induced as part of various stress responses. Two different pathways for synthesizing OLs are currently known: the OlsBA pathway first described in *Sinorhizobium meliloti*, and the OlsF pathway first described in *Serratia proteamaculans*. We identified in the genome of *Vibrio cholerae* O1 El Tor A1552 two genes encoding OlsF homologs, VC0489 is located on chromosome 1, whereas VCA0646 is located on chromosome 2. Both synthases, when expressed in *Escherichia coli*, caused the synthesis of OLs. Single mutants deficient in each of the OL synthases, double mutants deficient in both OL synthases, and mutants deficient in the transcriptional regulator PhoB were constructed and characterized. We corroborated that VC0489 is solely responsible for the synthesis of OLs under phosphate-limitation. The deletion of VC0489 reduced the growth velocity compared to the wildtype under phosphate-limiting conditions but not under phosphate-replete conditions. The expression of VCA0646 is favored under low salt growth conditions, and its deletion abrogates OL synthesis at low salinities. The absence of VCA0646 and, therefore, the lack of OLs under low salt conditions makes the respective mutant more susceptible to polymyxin than OL-forming strains. None of the mutants was affected in biofilm formation, swimming, or virulence assays using *Caenorhabditis elegans* or *Galleria mellonella*. Here, we describe two functional OL synthases present in a single bacterium for the first time, and we show evidence that OLs have an important function during the *V. cholerae* lifecycle.

## INTRODUCTION

Microorganisms frequently are exposed to changing environmental conditions. During the response of bacteria to abiotic stresses, the membrane has a vital function, with bacteria frequently adapting their membrane lipid composition (1). Phospholipids with a diacylglycerol backbone are the best-known membrane-forming lipids, but many other lipids can be present in bacterial membranes. Some of these membrane lipids are only formed by certain groups of bacteria or under specific stress conditions. One example of lipids whose synthesis and presence often are related to stress responses are ornithine lipids (OLs). OLs are phosphorus-free aminolipids found only in bacteria and absent from archaea or eukaryotes. In some bacteria, for example, *S. meliloti* or *Pseudomonas* sp., OLs are only formed under phosphate-limiting conditions (2–4), while in other bacteria, like many species of the genus *Burkholderia* or in *Rhizobium tropici* CIAT899, OLs are formed constitutively (5–8). The presence of OLs has also been related to increased resistance to elevated temperatures and low pH conditions (6, 8, 9), and OLs seem to have a function during the interactions between bacteria and eukaryotic hosts (5, 10). Recently, it was shown that ornithine lipid is a partial TLR4 agonist and NLRP3 activator (11).

Their basic structure is composed of a 3-hydroxy fatty acid linked by an amide bond to the α-amino group of ornithine and a second fatty acid linked by an ester bond to the 3-hydroxyl group of the first fatty acid (12, 13). These lipids can be formed by the acyltransferases OlsB and OlsA originally described in *S. meliloti* (14, 15) or by the bifunctional acyltransferase OlsF first described in *S. proteamaculans* (16). The OlsBA pathway is present in several α-and β-proteobacteria, a few γ-proteobacteria, and several actinomycetes. Genes encoding OlsF are present in a few γ-proteobacteria, δ-and ε-proteobacteria, and in bacteria belonging to the Cytophaga-Flavobacterium-Bacteroidetes (CFB) group. Based on the analysis of genomic DNA sequences, it has been estimated that about 50% of the sequenced bacterial species can form OLs, at least under specific growth conditions (12, 16). The presence of genes encoding OlsB or OlsF in bacterial genomes is a good indicator of their capacity to form OLs. For several bacteria whose genomes contain the respective genes, the presence of OLs was never described, and therefore, the roles of OLs and the conditions that favor their production have not been analyzed in depth in these organisms.

*V. cholerae* is a Gram-negative comma-shaped bacterium and the causative agent of the intestinal disease cholera. *V. cholerae* is found in brackish water and contaminated food. Outside of a human host, this bacterium can survive in bodies of water and marine animals, such as oysters and crabs. Even though *V. cholerae* is an extremely well-studied organism, the presence of OLs was not described until 2018, when Barbosa et al. (17) published that OLs are formed in the *V. cholerae* O1 El Tor strain N16961 under phosphate deprivation and that OL formation was dependent on the presence of VC0489 (OlsF) and a functional PhoBR two-component system.

In the present study, we describe that the *V. cholerae* O1 El Tor A1552 genome encodes two functional ornithine lipid synthase (OlsF) homologs. Mutants deficient in the OL synthase VC0489 show a reduced growth velocity and yield under phosphate-limiting conditions. The second OL synthase VCA0646 is transcribed in conditions of low to medium salinity, and OL accumulation is observed in *V. cholerae* under these conditions. Neither motility, biofilm formation, nor virulence of *V. cholerae* in *Caenorhabditis elegans* or *Galleria mellonella* were affected in the absence of OLs. On the other hand, the absence of OLs impacted the susceptibility of *V. cholerae* O1 El Tor to the effect of the cationic polypeptide antibiotic polymyxin B in low salinity conditions. This result suggests that the incorporation of OLs in the cell membrane of *V. cholerae* is not only a response to limiting phosphate concentrations but could also play an important role in the tolerance of this human pathogen to cationic microbicidals or other cell-membrane stressors, in reservoirs with low salinity, such as brackish or contaminated drinking water.

## RESULTS

### The *V. cholerae* genome encodes two functional OlsF homologs

Several years ago, we discovered the bifunctional ornithine lipid synthase OlsF in *S. proteamaculans* (16). Genes encoding OlsF-homologs were discovered in the genomes of a wide range of bacteria, including many species for which the formation of ornithine lipids (OLs) had not been described earlier, such as *V. cholerae*. In 2018, Barbosa et al. (17) described that OLs are formed in the *V. cholerae* O1 El Tor strain N16961 under phosphate deprivation and that OL formation was dependent on the presence of VC0489 (OlsF) and a functional PhoBR two-component system. Barbosa et al. expressed VC0489 heterologously in *E. coli* and observed that a PhoB-mutant could not induce OL synthesis under conditions of phosphate limitation. In their study, Barbosa et al. did not describe that the genomes of *V. cholerae* O1 El Tor strains, such as the reference strains N16961 and A1552 (18), encode two different OlsF homologs, VC0489 and, in addition, VCA0646 in chromosome 1 and chromosome 2, respectively. In our study strain, *V. cholerae* O1 El Tor A1552, VC0489 shows an identity of 33% and similarity of 48% on amino acid level with OlsF *from S. proteamaculans*. On the other hand, VCA0646 shows an identity of 30% and similarity of 46% on amino acid level with OlsF from *S. proteamaculans*. Both predicted *V. cholerae* proteins share an identity of 54% and similarity of 73% on amino acid level. VCA0646 is predicted to encode a 581 amino acid protein without transmembrane anchoring domains.

Both genes, VC0489 and VCA0646, were amplified by PCR, cloned into the expression vector pET17b, and expressed in *E. coli* BL21(DE3).pLysS. The resulting *E. coli* strains were labeled with [^14^C] acetate, and the lipids were extracted and separated by one-and two-dimensional thin-layer chromatography. In the vector control strain, only the phospholipids phosphatidylethanolamine (PE), phosphatidylglycerol (PG), and cardiolipin (CL) were detected (Fig. 1A), whereas in the two strains expressing a putative OL synthase, in addition to the phospholipids a lipid spot migrating as OL was detected (Fig. 1BC). The presence of OL in these lipid samples was also confirmed by LC-MS/MS (data not shown). Both OL synthases caused the formation of OLs in *E. coli*, which makes *V. cholerae* the first organism described with two functional OL synthases.

**Figure 1.**
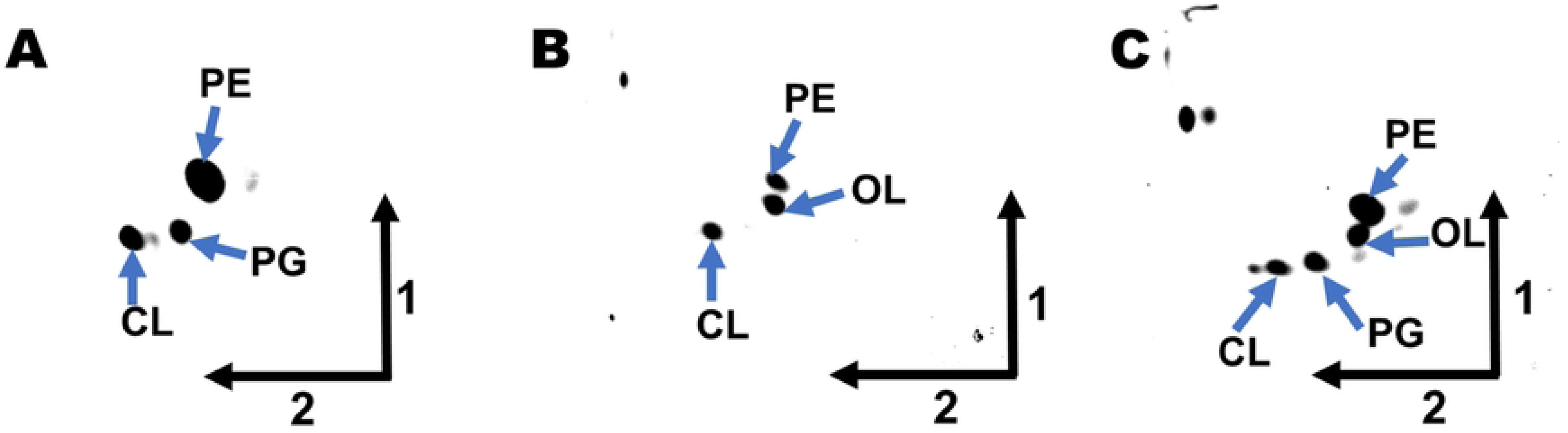
Ectopic expression of the putative ornithine lipid (OL) synthases VC0489 and VCA0646 from *V. cholerae* in *E. coli* BL21(DE3).pLysS promotes OL synthesis. Representative images of lipids labeled with [^14^C] acetate, extracted from *E. coli* strains harboring an empty control plasmid (A) or ectopically expressing VC0489 (B) or VCA0646 (C) and separated by thin-layer chromatography in two dimensions. Arrows indicate the first and the second dimension. PE: phosphatidylethanolamine; PG: phosphatidylglycerol; CL: cardiolipin; OL: unmodified ornithine lipid.

### VC0489-dependent OL synthesis activity contributes to optimal growth under phosphate-limiting conditions in *V. cholerae*

In some bacteria, for example, *S. meliloti*, *S. proteamaculans*, or *Pseudomonas* sp., OLs are only formed under phosphate-limiting conditions (2–4), while in other bacteria, like many species of the genera *Burkholderia* and *Agrobacterium* or in *Rhizobium tropici* CIAT899, OLs are formed constitutively (5–8). The reason for this discrepancy is not understood. *V. cholerae* formed the phospholipids PE, PG, and CL when grown in an LB medium or a standard M9 minimal medium with 64 mM phosphate but induced OL formation under conditions of phosphate limitation (17). Barbosa et al. showed that the OlsF homolog VC0489 is induced under these conditions and that transcription of the gene encoding VC0489 is controlled by the transcriptional regulator PhoB (17). The phenotype of a null mutant in VC0489 was not explored in their study (17); thus, we generated a deletion of VC0489 in *V. cholerae* A1552 (wildtype) and analyzed its growth in defined minimal medium (M9 medium) with phosphate concentrations from 1.28 mM to 64 mM. The latter corresponds to the standard phosphate concentration of this growth medium. A phosphate concentration of 4.2 mM was chosen for further experiments, representing a growth condition affecting growth without severely impeding it and, therefore, likely inducing the phosphate starvation response (Supplementary Fig. S1).

In addition, single deletion mutants were generated for VCA0646 and *phoB*, and also a double deletion mutant lacking VC0489 and VCA0646 was generated in the *V. cholerae* strain O1 El Tor A1552 (18). We also generated complementation constructs for VC0489 and VCA0646 in the expression plasmid pMMB67EH-Gen. We analyzed the growth of the wildtype strain and the single and double mutant strains in M9 growth medium with either 64 mM phosphate or 4.2 mM phosphate. In the absence of phosphate limitation (64 mM phosphate), all mutants grew as the wildtype (Fig. 2A). Under phosphate limitation (4.2 mM phosphate), the mutant strains lacking VC0489 or *phoB* showed reduced growth in comparison with the wildtype, whereas the single mutant lacking VCA0646 grew as the wildtype (Fig. 2B). The growth phenotype of the mutant strain lacking VC0489 in M9 media with 4.2 mM phosphate could be complemented when VC0489 was expressed from the pMMB67EH-Gen plasmid. We also observed a partial recovery of the growth-deficient phenotype in M9 media with 4.2 mM phosphate of the mutant strain lacking *phoB* when VC0489 was expressed from the P_tac_ promoter of the pMMB67EH-Gen plasmid. These observations strongly suggest that the product of VC0489 is required for optimal growth in conditions of phosphate limitation. Furthermore, our data also suggest that an important part of the growth defect that results from the absence of PhoB under phosphate-limitation growth conditions could be due to the inability to express VC0489.

**Figure 2.**
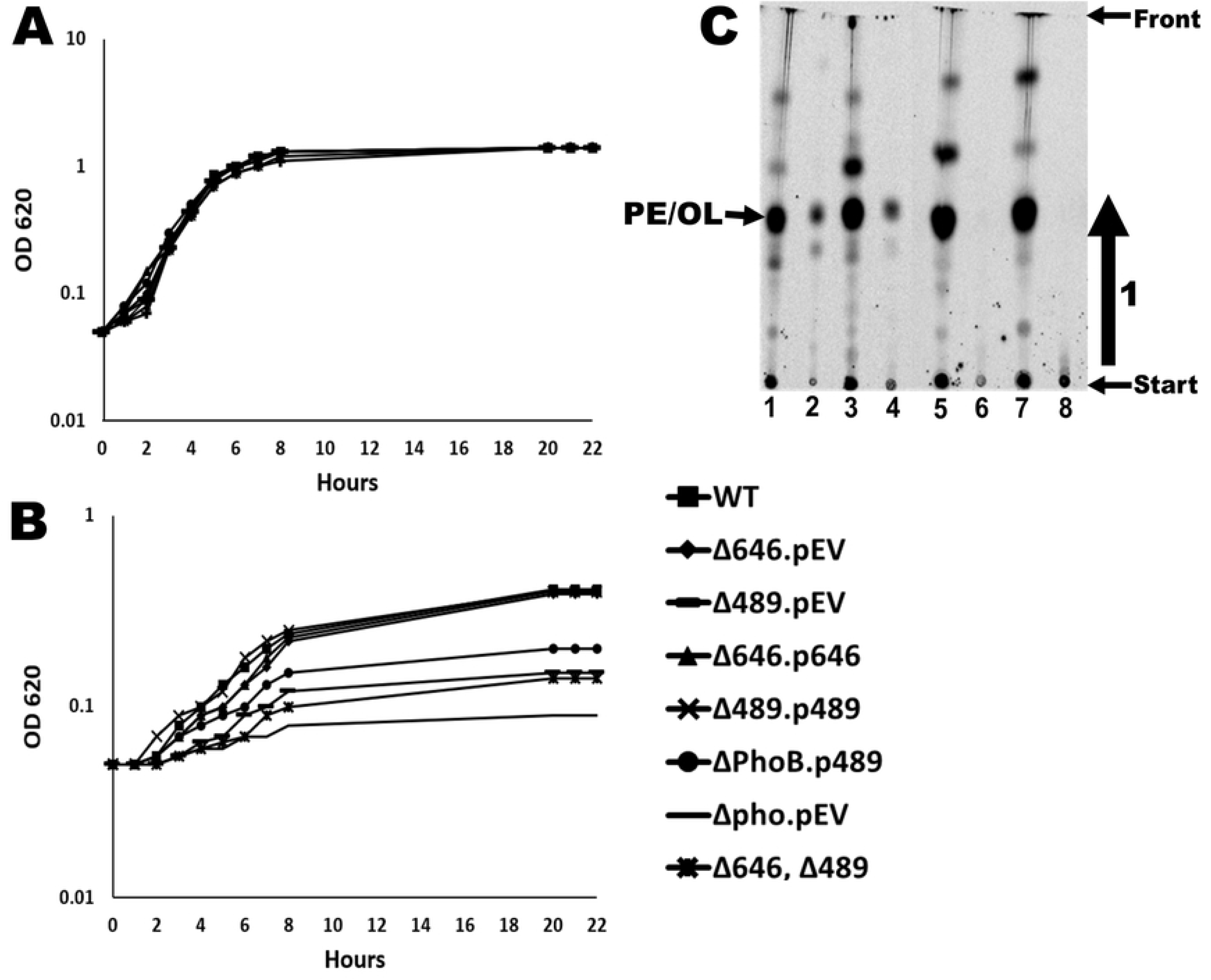
The absence of VC0489 results in reduced growth and lack of OL synthesis at limiting-phosphate growth conditions in *V. cholera*e. Growth curves of strains of interest in M9 medium with 64 mM phosphate (A) or in M9 medium with 4.2 mM phosphate (B). (C) Labeling with [^14^C]acetate or [^14^C]ornithine in M9 medium with 4.2 mM phosphate for 18 hours. Lane 1: *V. cholera*e wildtype [^14^C]acetate, lane 2: *V. cholera*e wildtype [^14^C]ornithine, lane 3: *V. cholera*e ΔVCA0646 [^14^C]acetate, lane 4: *V. cholera*e ΔVCA0646 [^14^C]ornithine, lane 5: *V. cholera*e ΔVC0489 [^14^C]acetate, lane 6: *V. cholera*e ΔVC0489 [^14^C]ornithine, lane 7: *V. cholera*e 2M (ΔVCA0646 and ΔVC0489) [^14^C]acetate, and lane 8: *V. cholera*e 2M (ΔVCA0646 and ΔVC0489) [^14^C]ornithine. The arrow in panel C indicates the direction of the chromatography. OLs and PE are not separated in this separation system, and their migration is indicated by an arrow.

Although a critical role for membrane lipid turnover under conditions of phosphate limitation has been suggested by different authors, this is, to our knowledge, the first time that a clear growth inhibition phenotype has been observed for a mutant deficient in OL synthesis under phosphate limitation.

To evaluate if the limited-phosphate growth conditions used in this study promote OL synthesis we labeled the lipids of the wildtype and mutant strains with [^14^C] acetate (to reveal all lipids) or with [^14^C] ornithine (to reveal almost specifically OLs). OLs can be detected under phosphate limitation in the wildtype and the mutant deficient in VCA0646, but not in the mutant deficient in VC0489 or the double mutant deficient in both genes (Fig. 2C). This result provides evidence that VC0489 promotes OL synthesis in M9 media with 4.2 mM phosphate, strongly suggesting that the lack of OL synthesis, in the mutant lacking VC0489, is responsible for the growth defect under phosphate limitation. Our data also indicates that VCA0646 does not contribute to OL formation under the phosphate-limiting growth conditions used in this study (17).

### VCA0646 is expressed at low to medium salinity and is responsible for OL formation under these conditions

We have shown that VCA0646 has OL synthase activity when expressed in a heterologous host (Fig. 1C). However, its contribution to OL formation was not detectable under low phosphate conditions in *V. cholerae* (Fig. 2C, lane 6). Thus, we next asked in which conditions this OL synthase might be expressed and promote OL synthesis.

Several members of the *Vibrio* genus are halophiles, a characteristic that limits their ability to colonize particular niches. Pandemic *V. cholerae* stains, however, can grow in a relatively broad range of salinity concentrations, a characteristic that allows them to survive in brackish water and freshwater (19). We speculated that changes in salinity might require adaptations in the cell membrane properties, which could involve the formation of OLs.

We first analyzed if changes in NaCl concentration affected the production of OLs in *V. cholerae* cells grown in LB broth. Since the typical concentration of NaCl used for the growth of *V. cholerae* in LB broth is about 0.17 M, we evaluated OL synthesis in LB broth with 0.0375, 0.075, 0.1, and 0.2 M NaCl using the wildtype and single mutants deficient in one or the other OL synthase harboring an empty or a complementing plasmid. After labeling growing cells with [^14^C]ornithine for 18 hours, we observed accumulation of OLs in the wildtype strain and the single mutant strain lacking VC0489 in cells grown in LB with 0.0375, 0.075 and 0.1 M NaCl but not at 0.2M NaCl (Fig. 3). This OL synthesis depended on the presence of VCA0646, a mutant strain lacking this gene could not accumulate OLs under these conditions but could be complemented in trans.

**Figure 3.**
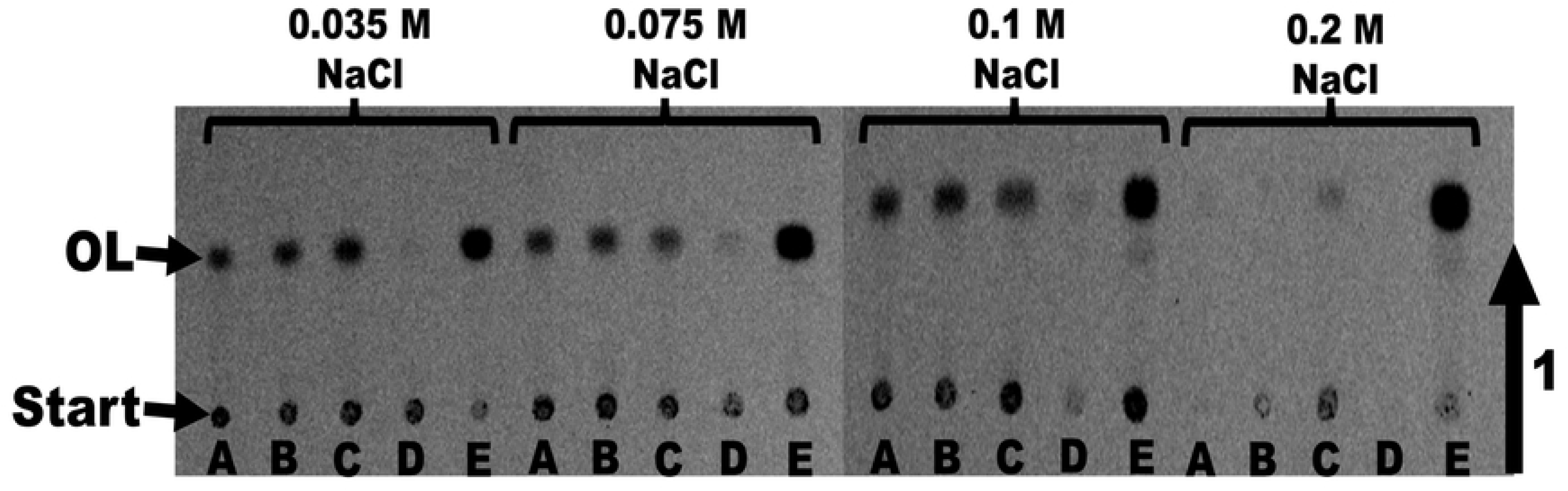
VCA0646 is responsible for OL synthesis in *V. cholerae* under low salt conditions. Labeling with [^14^C]ornithine in LB medium with different salt concentrations (0.0375 M NaCl, 0.075 M NaCl, 0.1 M NaCl, and 0.2 M NaCl) for 18 hours. *V. cholerae* wildtype (A), mutant ΔVC0489 harboring an empty plasmid (B), mutant ΔVC0489 harboring the complementing plasmid (C), mutant ΔVCA0646 harboring an empty plasmid (D), and mutant ΔVCA0646 harboring the complementing plasmid were used. Lipids were extracted according to Bligh and Dyer and separated by one-dimensional TLC. Radioactivity can be detected in the origin, where the samples are applied and in OL.

The promoter controlling the expression of VCA0646 or VC0489 in the plasmid is constitutive, explaining why we observe OL formation at 0.2M NaCl in conditions where the wildtype does not form OLs. These results strongly suggest that the product of VCA0646 is responsible for OLs production in rich media at medium to low salinity (Fig. 3). The growth of the mutant strains that were not able to synthesize OLs was not significantly affected in LB medium with reduced NaCl (Supplementary Fig. S2). This would suggest that the formation of OLs is induced under these conditions but that they are not required for growth at low salinities, at least under the conditions tested.

The requirement of VCA0646 for OLs synthesis at low salinities suggests that the expression of this gene might respond to changes in salinity. VCA0646 is the last gene in a putative operon encoded in the complementary strand and composed of 4 genes (VCA0649 to VC0646) (Figure 4A). To analyze the expression of VCA0646, we first evaluated whether this gene had its own promoter or if its expression could come from a promoter upstream of VCA0649. We generated plasmid constructs with transcriptional fusions with upstream sequence from VCA0649 (P_VCA0649_-*lux*) or VCA0646 (P_VCA0646_-*lux*) fused to the reporter genes *luxCDABE* that encode a bacterial luciferase and its substrate. The transcriptional fusion P_VCA0649_-*lux* showed lower activity levels in all conditions and was not induced under reduced salt concentration (Figure 4B). On the other hand, the transcriptional fusion P_VCA0646_-*lux* was induced when salt concentrations were reduced (Figure 4C).

**Figure 4:**
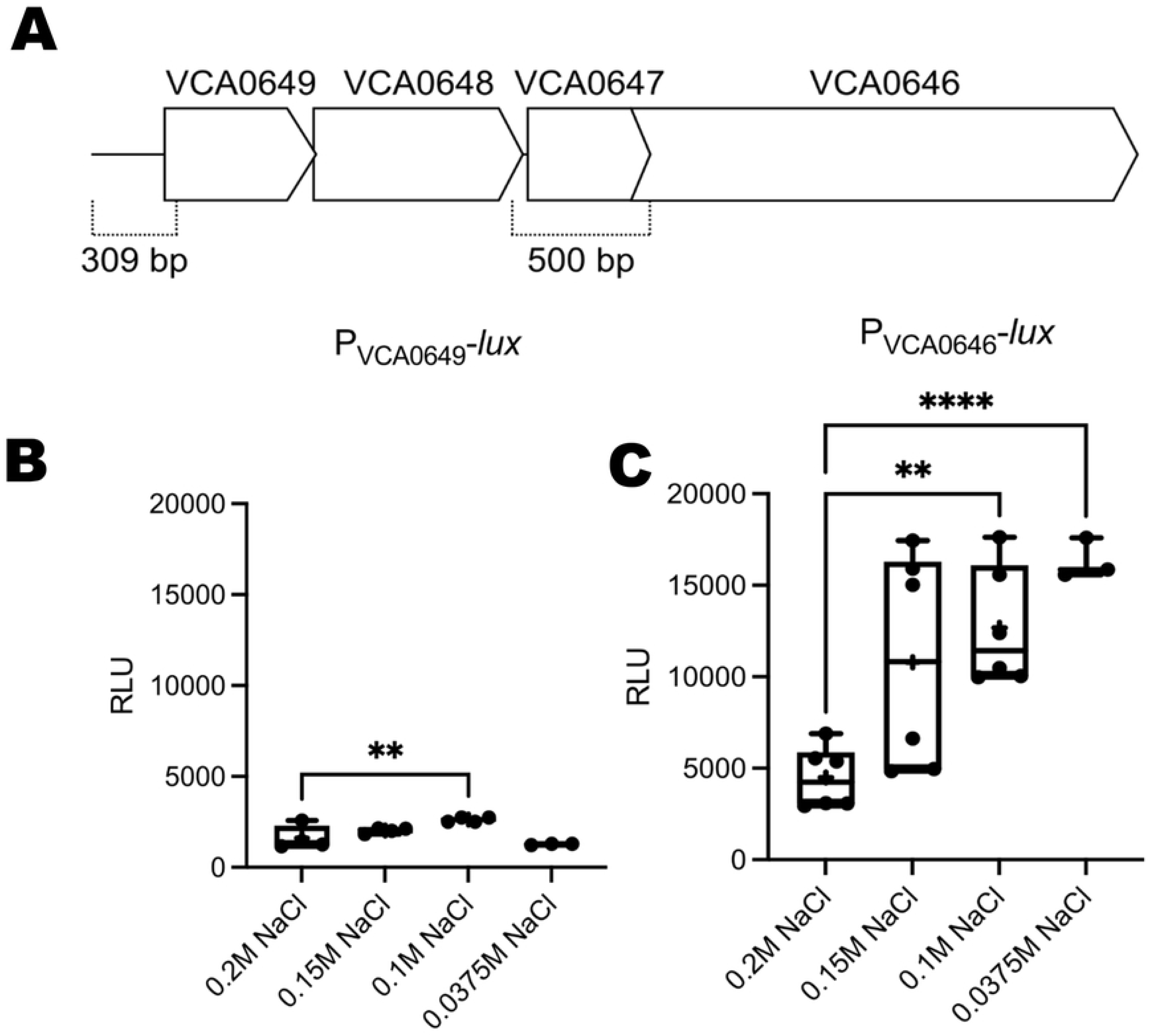
The PVCA0646 promoter has a higher expression at lower salinities in *V. cholerae*. (A) Illustration depicting the genetic arrangement of the gene cluster VCA0649 to VCA0646. The location of the regions used to generate transcriptional fusions with the reporter genes *luxCDABE* is indicated with dotted lines with their respective length shown below. (B) and (C) Box plots representing the relative luminescence units obtained from cells harboring either a P_VCA0649_-*luxCDABE* or a P_VCA0646_-*luxCDABE* transcriptional fusion grown in different NaCl concentrations. Experiments were done twice, and at least three biological replicates were analyzed. A Brown-Forsythe ANOVA test was used to evaluate statistically significant differences among means. This was followed by a Dunetts T3 multiple comparison test to compare between specific means. Adjusted P values below 0.05 were considered significant. Adjusted P values ** ≤ 0.01; **** ≤ 0.0001

This result suggests that there is an active promoter immediately upstream of VCA0646. More in-depth analysis would be required to characterize these two promoters and the presence and sequence identity of potential polycistrons within this region. Our results also revealed that the activity of the P_VCA0646_ promoter is higher at lower concentrations of NaCl. This is more evident when comparing growth conditions in the presence of 0.2 M NaCl versus 0.1 M or 0.0375 M NaCl. Together, our results suggest that the expression of the OL synthase VCA0646 is induced under low NaCl concentrations and, therefore, is responsible for synthesizing OLs in *V. cholerae* under these conditions.

### The absence of OLs affects resistance to polymyxin but does not affect biofilm formation, motility, or virulence in *C. elegans* and *G. mellonella*

We next wanted to find out if the presence of OLs in *V. cholerae* is important for virulence or virulence-related traits such as motility, biofilm formation or polymyxin resistance. All four strains were equally susceptible to polymyxin B when grown in LB medium with 0.2 M NaCl (Fig. 5A). We observed reduced susceptibility to polymyxin B, in the wildtype strain and the mutant strain ΔVC0489 but not in the strains that lack VCA0646, when grown in the presence of 0.0375 M NaCl when compared to conditions of growth in LB medium with 0.2 M NaCl (Fig. 5A). When we complemented the mutant strain ΔVCA0646 with a copy of VCA0646 in trans, its polymyxin resistance returned to wildtype level. Since VCA0646 is required for OL formation under low-salinity growth conditions, our results suggest that the synthesis of these lipids contributes to changes in the susceptibility to polymyxin B. OLs have been described to be present in the outer membrane in some proteobacteria, where polymyxin interacts with Lipid A. The increased resistance to polymyxin B may be due to the accumulation of the zwitterionic lipid OL in the outer membrane, which could dilute the negative charges of the lipid A molecules, thereby making the OL-forming strains more resistant.

**Figure 5.**
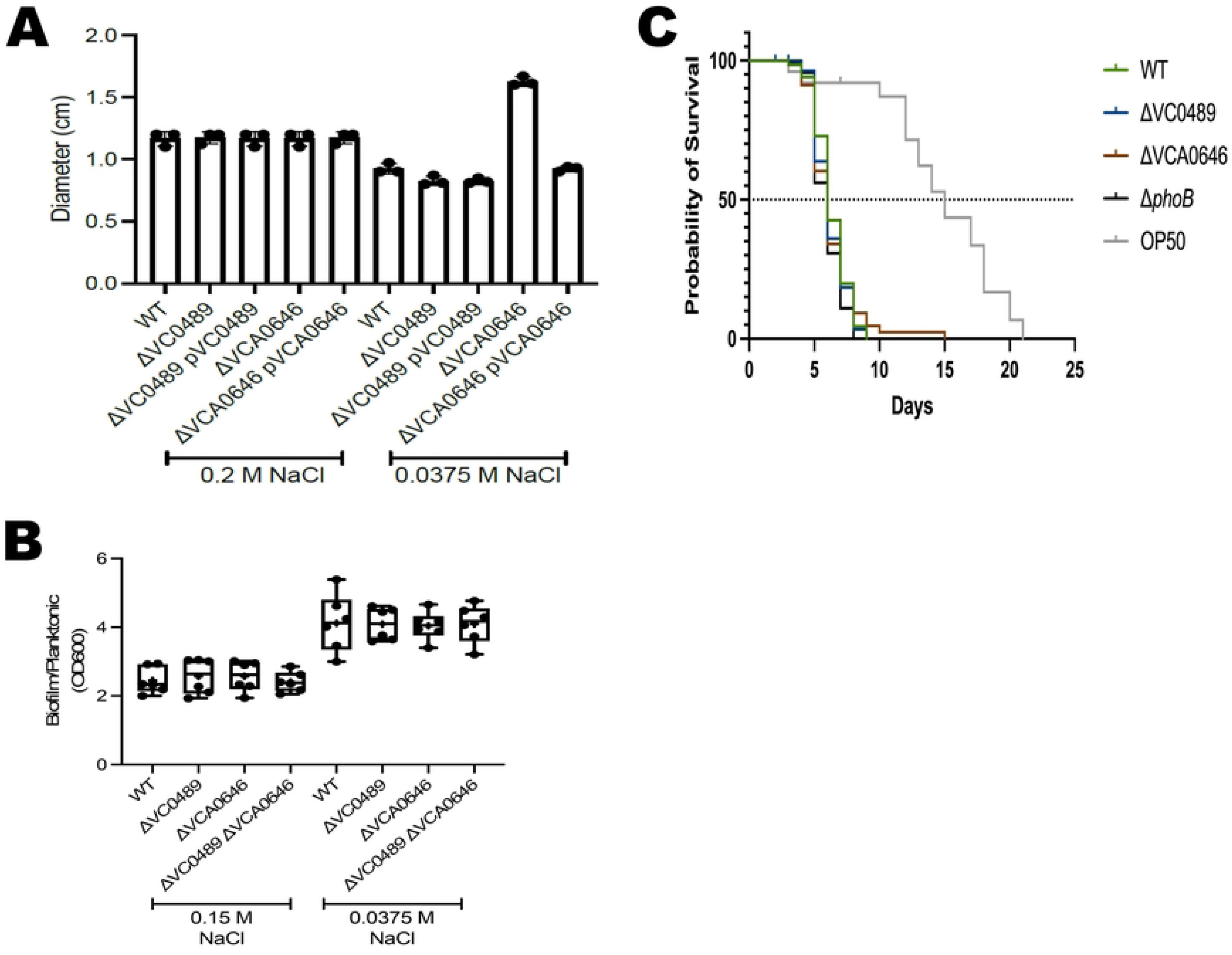
The presence of OLs affects resistance to polymyxin but does not affect swimming motility, biofilm formation, or virulence. (A) Bar graph showing the mean and standard deviation of the diameter of polymyxin B-dependent sensitivity halos. The indicated strains were grown in LB medium with 0.2 M NaCl (left) or LB medium with 0.0375 M NaCl (right). The absence of VCA0646 under low salt conditions makes the cells susceptible to polymyxin B. (B) Box plots representing the measurements obtained from the quantification of biofilms grown for 9 hours over a PVC surface relative to planktonic growth in LB medium containing either 0.15 M NaCl or 0.0375 M NaCl. (C) Survival curves show the kinetics of *Caenorhabditis elegans* killing by different bacterial strains. The *V. cholerae* data are compiled from three independent assays. The OP50 trace corresponds to a representative experiment. *p*<0.0001, OP50 vs *V. cholerae* strains.

We next compared the ability of the wildtype strain and the single and double mutants in genes VC0489 and VCA0646, to form biofilms under static conditions over a polyvinyl chloride (PVC) surface in LB media with 0.15 M NaCl or 0.0375 M NaCl. All strains showed a slight increase in biofilm formation when grown in the presence of 0.0375 M NaCl when compared to growth in 0.15 M NaCl, but we did not observe differences in biofilm formation capability between these strains under the conditions tested (Fig. 5B). We also assayed the swimming motility of the mutant strains but no differences were observed (data not shown).

We then evaluated if the absence of the genes required for OL biosynthesis, or their regulator, *phoB*, affects the ability of *V. cholerae* to kill the nematode *Caenorhabditis elegans*. As reported previously (20), worms fed with the wildtype *V. cholerae* strain die faster than the ones fed with *E. coli* OP50, the standard nematode food (median survival = 6 days for *V. cholerae* A1552 vs 15 days for *E. coli* OP50). We did not observe a significant difference in the killing capability when comparing the *V. cholerae* wildtype strain with the mutants lacking each of the *olsF* genes (ΔVC0489 and ΔVCA0646) or *phoB* in three independent assays (Fig. 5C).

The virulence of the *V. cholerae* strains on *G. mellonella* larvae was also evaluated. No differences in mortality were observed between the larvae injected with the wildtype or the different mutants (data not shown).

## DISCUSSION

OLs are phosphorus-free membrane lipids that can be formed by about 50% of the sequenced bacterial species but are absent from archaea and eukaryotes (1, 21). Two different pathways for OL synthesis are known: the OlsBA pathway, first described in the alpha-proteobacterium *S. meliloti*, and the OlsF pathway, first described in the gamma-proteobacterium *S. proteamaculans* (1, 14–16). Here, we report on the characterization of two OL synthases belonging to the OlsF family in *V. cholerae* called VC0489 and VCA0646. One is located on chromosome 1, and the other on chromosome 2. We found that both synthases can produce OL in the heterologous host *E. coli* when overexpressed using an IPTG inducible system. This is the first report showing that VCA0646 is an active OlsF homolog and it is the first report of the presence of two functional OlsF homologs in a single organism. We also found that both OlsF homologs can promote OL synthesis in *V. cholerae,* albeit under different growth conditions. VC0489 is responsible for OL synthesis during growth under phosphate limitation, while VCA0646 synthesizes OL in media with low to medium salinity. The absence of VC0489 or VCA0646 causes growth sensitivity phenotypes in media with low phosphate or in the presence of the antimicrobial polymyxin B, respectively. This strongly suggests that the synthesis of OL plays a role in the adaptation of *V. cholerae* to certain stressful conditions that could potentially compromise the cell membrane integrity and/or function, especially in reservoirs with low salinity, such as brackish or contaminated drinking water.

When searching bacterial genomes for genes encoding OlsB or OlsF, one usually identifies one or the other. To our knowledge, *V. cholerae*, closely related species, such as *V. paracholerae* and *V. terriae*, but also *V. vulnificus* have two genes encoding OlsF homologs. Our result further revealed that both OlsF homologs from *V. cholerae* are functional OL synthases implying an important function of OL in this organism, possibly during survival under low phosphate conditions or brackish water. VC0489 is mainly expressed in response to phosphate limitation (17). Under this condition, resources for phospholipid synthesis are scarce, and 40% or more of the membrane lipids are OLs. This could have important implications in multiple cellular processes, including maintenance of active division, assembly of cellular appendages and adhesins, and, in consequence, biofilm formation, motility, and bacterial competition or pathogenic capability. Future studies will be needed to evaluate in an integrated manner how OLs contribute to these processes.

In this study, VCA0646 was found to contribute to OL synthesis in media with medium to low concentrations of sodium salts. Under this condition, the cell membrane is composed of phospholipids and phosphorus-free OL. Our results suggest that in rich media with low salinity, the presence of OL is not crucial for processes such as biofilm formation and swimming motility. We cannot discard yet that under different conditions or with experimental approaches with single-cell resolution we could detect phenotypic effects for the absence of OL synthesis for these important cellular processes. However, we did observe differences in polymyxin B sensitivity that could be associated with the presence of OL in the cell membrane. Cells grown at high salinity (0.2 M) were more sensitive to polymyxin B than cells grown at low salinity (0.0375 M). Our results strongly suggest that the sensitivity phenotype is mostly attributable to the lack of OL synthesis since growth in 0.2 M NaCl-containing media severely inhibits OL synthesis. Furthermore, the higher tolerance to polymyxin B antimicrobial properties of cells grown in media with 0.0375 M NaCl is lost in the absence of VCA0646 which we also showed was necessary for OL synthesis under this growth condition. The negatively charged phosphate groups in the lipid A present in the outer leaflet of the outer membrane are targets of cationic peptides such as polymyxin B. Polymyxin B replaces Ca^2+^ and Mg^2+^ ions that usually help to stabilize the outer membrane and make it thereby more permeable (22). OL has been shown to accumulate in different Gram-negative bacteria in the outer membrane (8) and we speculate that the accumulation of the zwitterionic OL in the outer membrane acts as a shield to protect the anionic phosphate groups.

Mechanisms for polymyxin B resistance that have been reported before involve the removal of phosphate groups or a chemical modification of the phosphate groups with cationic groups such as ethanolamine or aminoarabinose (23, 24). In *V. cholerae* O1 El Tor, the most well-understood polymyxin B resistance mechanism involves the modification of lipids with glycine or diglycine amino acid residues by the AlmE, AlmF, and AlmG (25). The genes that encode this lipid A modification system are regulated by the CarRS two-component system (26). More recently, yet another lipid A modification system was uncovered in *V. cholerae* O1 El Tor, which involves an EptA ortholog that can transfer phosphoethanolamine (pEtN) to lipid A under conditions of low pH (27). Our results suggest that polymyxin B tolerance can also be favored by the modification of the lipid composition of the cell membrane under particular growth conditions, adding a further level of protection against this antimicrobial compound and perhaps other cationic peptides. While AlmE, which is part of the glycine-diglycine modification system, has been shown to affect biofilm formation in two different strains of *V. cholerae* O1 El Tor strains (26), the contribution of the pEtN modification to biofilm formation has not been investigated, and our results did not reveal an effect of OL synthesis on this important social behavior. The deletion of the operon that encodes the glycine-diglycine modification system (*almEFG*) did not affect colonization of the mouse small intestine, while the deletion of their regulator CarR does, in a strain-specific manner. This would suggest that the lack of effect on small-intestine colonization of the deletion of *almEFG* might not be due to lack of expression. Perhaps the presence of the pEtN modification system and OL synthesis could contribute to resistance to cationic peptides such as the cathelin-related antimicrobial peptide, which is produced in the small intestine of neonatal mice (28).

Alterations in cell membrane lipid composition or lipopolysaccharide modifications could potentially contribute to the fitness of environmental *Vibrio* strains with pandemic capabilities. Altogether, our results further support the premise that cell membrane remodeling plays an important role in the physiology of the human pathogen *V. cholerae* O1 El Tor. More studies will be needed to evaluate individual contributions of cell-membrane modifying systems and their connection to improved adaptability and pathogenic potential.

## MATERIALS AND METHODS

### Bacterial strains and growth conditions

This study used *V. cholerae* O1 El Tor A1552 [15] as wildtype strain (Table 2). *E. coli* DH5α (29) was used as a host strain for molecular biology experiments, and *E. coli* BL21(DE3).pLysS (30) was used as a host strain for the overexpression of the two putative OL synthases VC0489 and VCA0646.

Bacteria were grown in Lysogeny broth (LB; 5 g yeast extract, 10 g peptone, 10 g NaCl per liter and adding 1.5 % agar (w/v) for solid medium) or in M9 minimal medium designed for growth under phosphate-limiting conditions. The NaCl concentration of the standard LB medium is 0.17 M, which was varied for some experiments as indicated throughout the text. Standard M9 medium was prepared as a 5x salt solution (56.3 g Na_2_HPO_4_ x 7H_2_0, 15 g KH_2_PO4, 2.5 g NaCl and 5 g NH_4_Cl per liter). After autoclaving, MgSO_4_ was added to a final concentration of 2 mM, glucose to a final concentration of 20 mM, CaCl_2_ to a final concentration of 100 µM and MEM vitamin solution (Thermo Fisher, product #1120052) to a final concentration of 1x. In some growth experiments, the phosphate concentration, which is 64 mM under standard conditions, was reduced to 12.8 mM, 6.4 mM, 4.2 mM, 2.56 mM, or 1.28 mM. A final phosphate concentration of 4.2 mM was used for most experiments under phosphate limitation.

### Generation of genetic constructs for deletion, overexpression and reporter-based transcription analysis. fusions

Oligonucleotide primers used to construct deletions and transcriptional fusions are listed in Table 1. Primers VC0489_01 and VC0489_02 were used in a PCR to amplify about 1.0 kb of genomic DNA upstream of the VC0489 gene from *V. cholerae* O1 El Tor A1552, introducing EcoRI, SalI, and BamHI sites into the PCR product. Similarly, primers VC0489_03 and VC0489_04 were used to amplify about 1.0 kb of genomic DNA downstream of the VC0489 gene from *V. cholerae* O1 El Tor A1552, introducing BamHI and XbaI sites into the PCR product. After digestion with the respective enzymes, PCR products were cloned as EcoRI/BamHI or BamHI/XbaI fragments into pUC19 to yield the plasmids pUCVC01 and pUCVC02, respectively. Then, the BamHI/XbaI fragment from pUCVC02 was subcloned into pUCVC01 to yield pUCVC03. Plasmid pUCVC03 was digested with SalI and XbaI to subclone the regions usually flanking the *V. cholerae* O1 El Tor A1552 VC0489 gene into the suicide vector pGP704-sacB (31) to yield pUCVC04. Via diparental mating using *E. coli* S17-1 (32) as a mobilizing strain, pPMAV04 was introduced into the wildtype strain *V. cholerae* O1 El Tor A1552. The genetic constructs used to delete VCA0646 and *phoB* were assembled into the suicide plasmid pGP704-sacB through isothermal assembly (“Gibson assembly”). This plasmid has an R6K origin of replication and an origin of transfer (oriT). The oligonucleotides used to amplify the assembly parts were designed using the NEBuilder Assembly tool (https://nebuilder.neb.com). All primers used are described in Table 1. Plasmid pGP704-sacB was linearized through inverse PCR. The homologous recombination substrates consist of 500 bp of upstream and downstream sequences flanking the deletion targets and were amplified with oligonucleotides that allow assembling them together and to the linearized pGP704-sacB. The amplification products were purified and concentrated using the DNA Clean and Concentrator Kit from Zymo Research. The isothermal assembly was performed using the NEBuilder HiFi DNA Assembly Master Mix. The assembly reaction was set on ice using 50 to 100 ng of each assembly part and 2 μl of the NEBuilder HiFi DNA Assembly Master Mix in a final volume of 6 μl. The assembly reaction was incubated 1 hour at 50 °C and then used to transform chemically competent cells of the *E. coli* strain SM10λ*pir* or S17λ*pir* following standard procedures. Positive clones were selected after genotyping through colony PCR using oligonucleotides that flank the assembly site. Sanger sequencing was used to verify the fidelity of amplification and assembly.

**Table 1.**
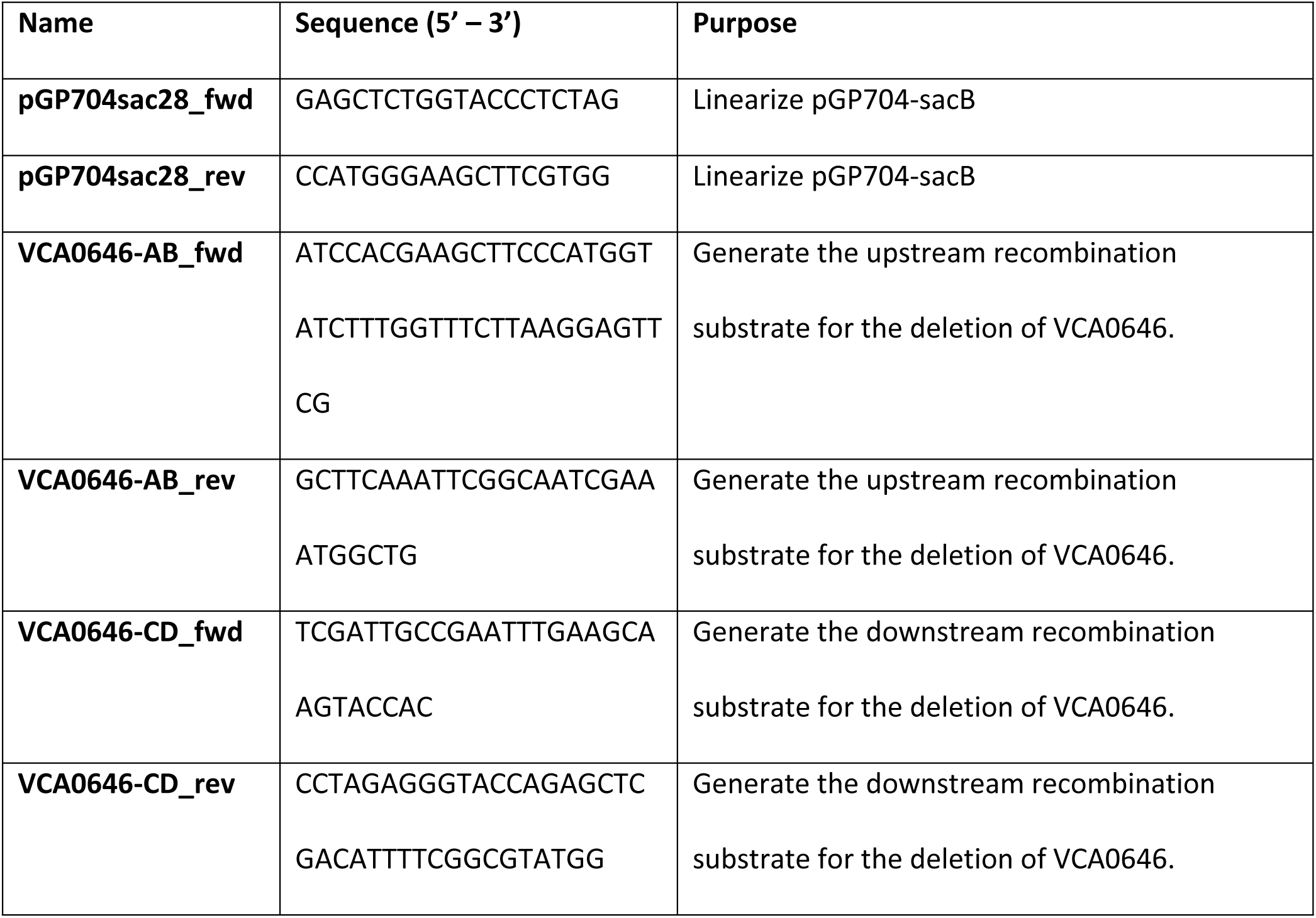
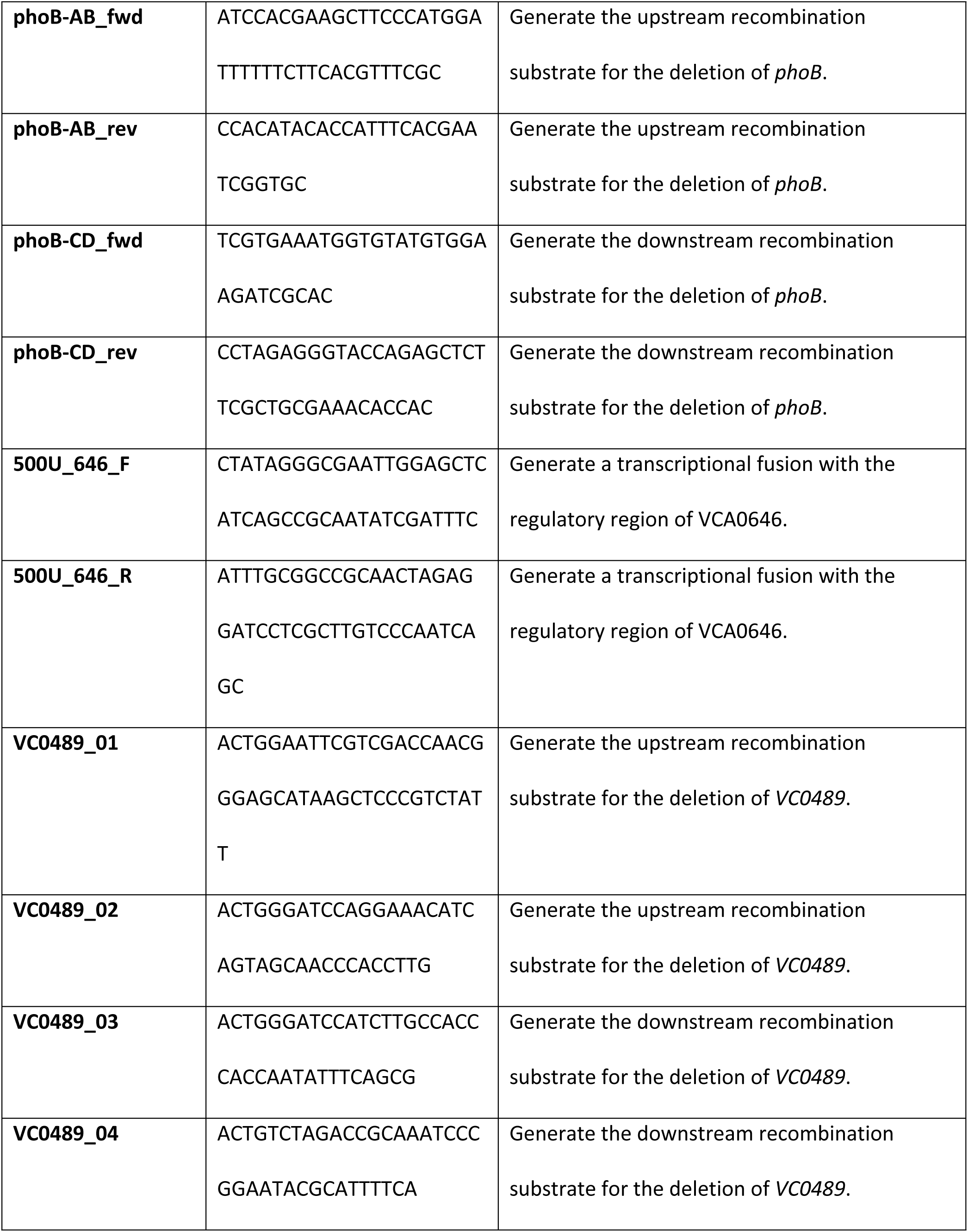
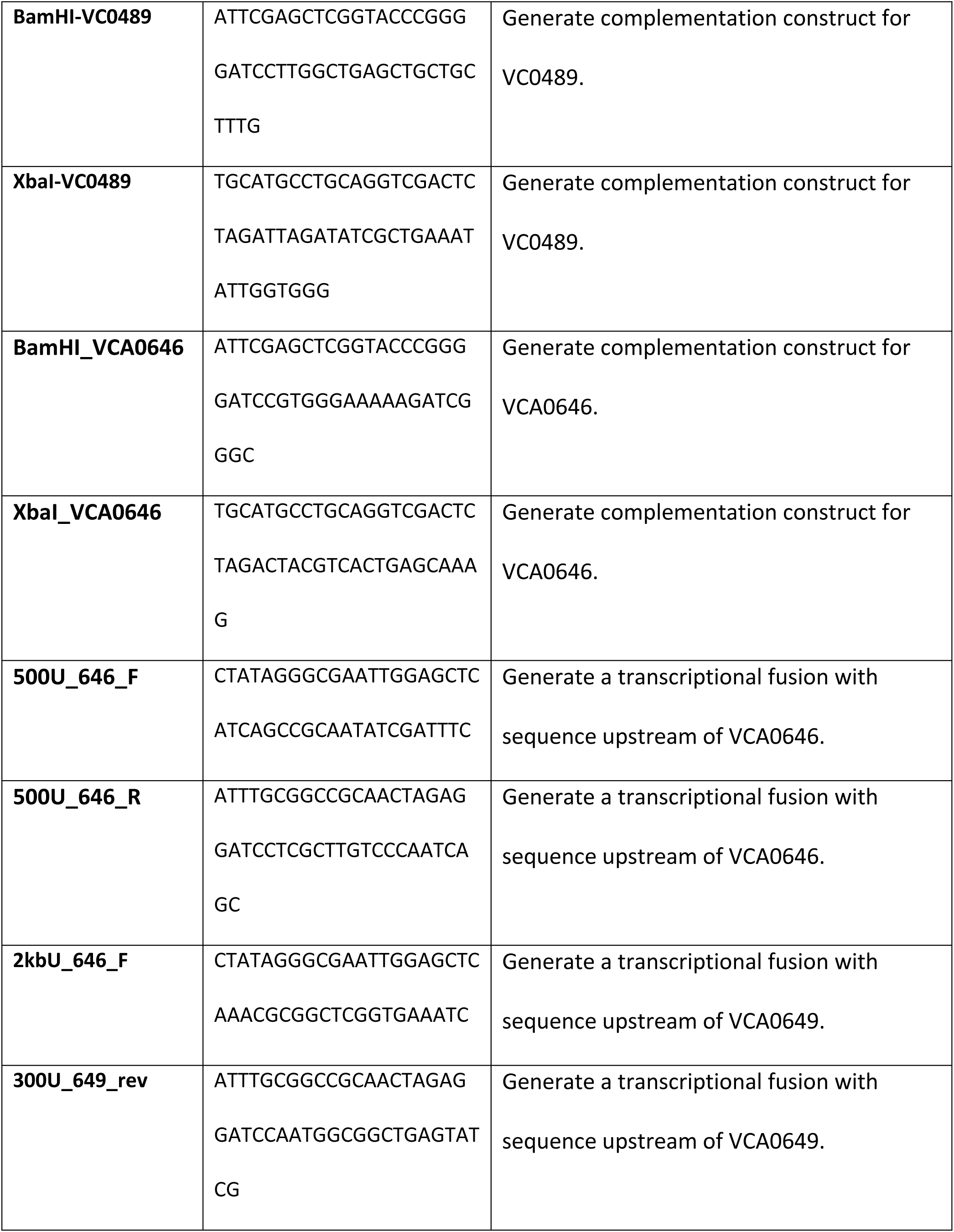
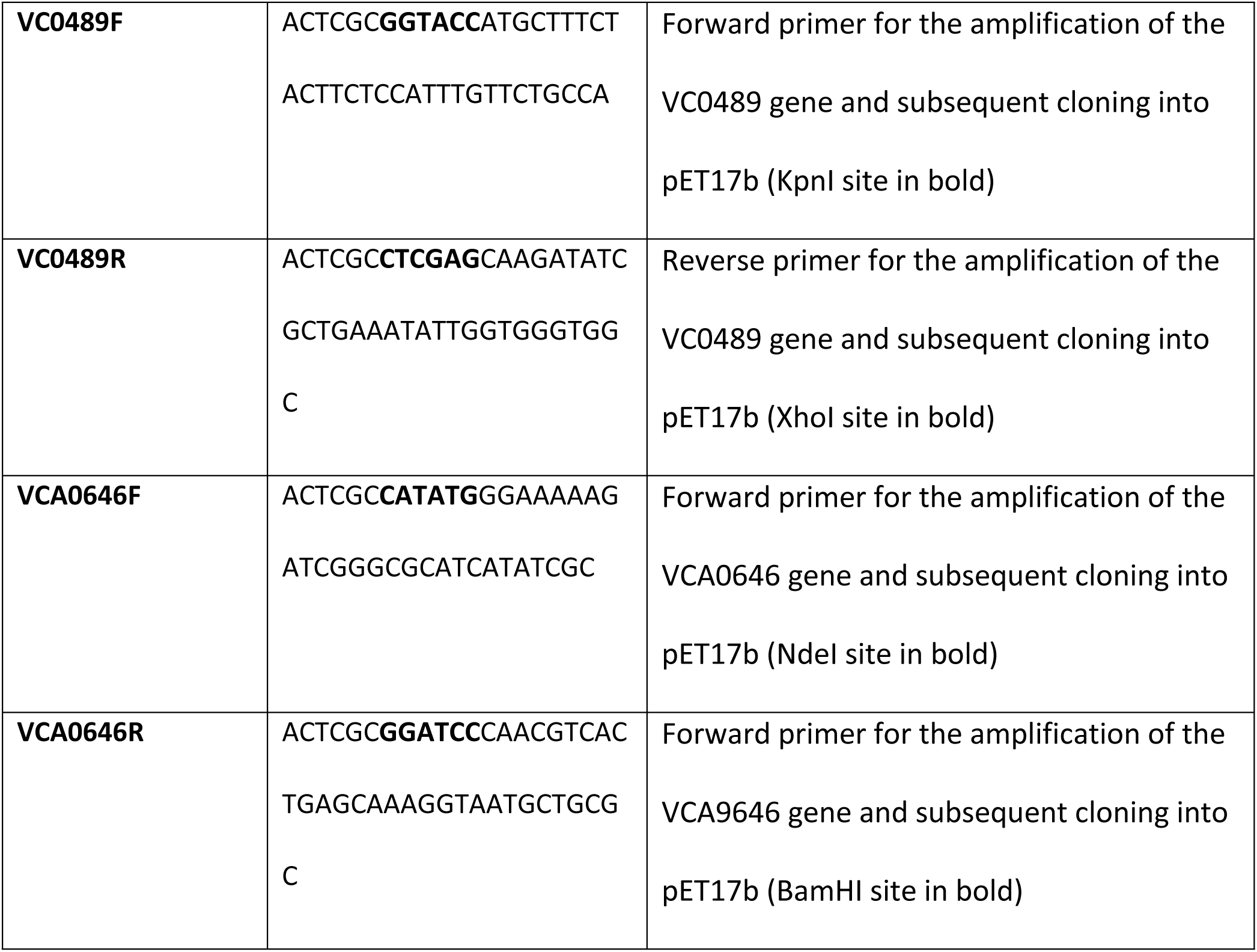
Oligonucleotide primers used in this study. All sequences are written in 5’to 3’direction

**Table 2.** Plasmids used in this study.

| Name | Purpose | Reference |
| --- | --- | --- |
| pDZ_4 | Deletion of VCA0646 | This study |
| pDZ_5 | Deletion of *phoB* | This study |
| pDZ_59 | Complementation with VC0489 | This study |
| pDZ_57 | Complementation with VCA0646 | This study |
| pDZ_55 | Transcriptional fusion PVCA0646-<br>*luxCDABE* | This study |
| pDZ_58 | Transcriptional fusion PVCA0649-<br>*luxCDABE* | This study |
| pUC19 | Cloning vector, carbenicillin<br>resistant | (39) |
| pGP704-*sacB* | Suicide vector, carbenicillin-<br>resistant | (31) |
| pUCVC01 | 1 kb fragment upstream of<br>VC0489, cloned as EcoRI,<br>Sall/BamHI fragment in pUC19 | This study |
| pUCVC02 | 1 kb fragment downstream of<br>VC0489, cloned as BamHI /XbaI<br>fragment in pUC19 | This study |
| pUCVC03 | 1 kb upstream and 1 kb<br>downstream sequences flanking<br>VC0489 cloned into pUC19 | This study |
| pUCVC04 | Suicide vector for construction of<br>mutant of VC0489. | This study |
| pMMB67EH-Gen<br>(pFY_1554) | Expression plasmid for use in *V.*<br>*cholerae* | Samuel Miller and Fitnat Yildiz |
| pET17b | Expression plasmid for use in *E. coli*<br>BL21(DE3).pLysS | (30) |
| pET17b.489 | pET17b-with gene VC0489 cloned<br>as KpnI/XhoI-fragment | This study |
| pET17b.646 | pET17b-with gene VCA0646 cloned<br>as NdeI/BamHI-fragment | This study |

The deletion constructs were mobilized to *V. cholerae* A1552 through biparental mating, using as donor *E. coli* strains SM10λ*pir* or S17λ*pir*. Donor and recipient strains were mixed in a 1:1 proportion (in a final volume of 2 ml) and centrifuged at 4000 x g for 3 minutes. The pellet was suspended in 150 μl of LB, 50 μl of the suspension was transferred in triplicate to an LB agar plate and incubated overnight at 37 °C. The mating spots were suspended in 1 ml of LB and diluted 1:10 in fresh LB. The diluted samples were spread over LB agar plates supplemented with 100 μg/ml rifampicin and 100 μg/ml ampicillin or carbenicillin to select for single homologous recombination events that result in plasmid integration. Single recombinants were streaked over LB agar plates with no antibiotics or selective TCBS agar plates. Single colonies were used to prepare cultures in LB broth. Cultures were grown for 6 to 8 hours in LB broth in the absence of antibiotic selection and then used to streak over LB agar plates with no NaCl and supplemented with 10% sucrose. Double homologous recombination results in the loss of the counterselection gene *sacB,* which confers sensitivity to the presence of sucrose in the growth media. Putative double homologous recombinants were patched over TCBS or LB agar plates and replica plated on LB agar plates supplemented with 100 μg/ml carbenicillin/ampicillin. The gene VC0489 was deleted through allelic exchange with a spectinomycin resistance cassette, double recombinants were also patched in LB agar plates supplemented with 100 μg/ml spectinomycin. Double homologous recombinants should be Amp^S^, and in the case of ΔVC0489 Spe^R^. To identify double recombinants that lack the genes of interest, we conducted colony PCR using primers that flank the structural gene. In the case of the construction of the mutant deficient in VC0489, candidates were grown, and their lipid composition was analyzed. One strain presented the expected phenotype which is the absence of the OL (S1).

The complementation constructs were assembled on a pMMB67EH-Gen plasmid through restriction digest and Gibson Assembly. The pMMB67EH-Gen plasmid was linearized using BamHI-Hf and XbaI restriction enzymes. The pairs of primers used to amplify VC0489 or VCA0646 were designed using the NEBuilder Assembly tool (https://nebuilder.neb.com) to generate complementary ends that enable assembly into the linearized pMMB67EH-Gen plasmid through Gibson assembly as described above. S17λ*pir* chemically competent cells were transformed with the assembly mix and selected over LB-agar with Gentamicin 15 μg/ml. Positive clones were identified through colony PCR using oligonucleotides that flank the assembly site. The fidelity of the assembled construct was verified with Sanger sequencing.

The transcriptional fusions of the regulatory region of VCA0649 or VCA0646 and the reporter genes *luxCDABE* were generated through Gibson assembly following the same protocol described above. A pair of oligonucleotides was designed, using the NEBuilder Assembly tool, to amplify 258 bp upstream of the coding sequence of VCA0649 and 51 bp of its coding sequence; or 428 bp upstream of the coding sequence of VCA0646 and 72 bp of its coding sequence. These oligos allow the assembly of the amplification product into the pBBRlux plasmid previously linearized with the restriction enzymes SacI-HF and BamHI-HF. The assembly reactions were used to transform S17λ*pir* chemically competent cells selecting over LB-agar plates with Chloramphenicol 20 μg/ml. Positive clones were identified through colony PCR using oligonucleotides that flank the assembly site. Sanger sequencing was used to verify the fidelity of amplification and assembly.

### Lipid analysis of bacterial strains

The lipid compositions of bacterial strains were determined following labeling with [1-^14^C] acetate or [^14^C] ornithine (ARC, American Radiochemicals). Cultures (1 ml) of *E. coli* BL21(DE3).pLysS with different expression plasmids were grown, and at an OD_620_ of 0.3 to 0.4, 0.2 mM IPTG, and 1 µCi of [^14^C]acetate (ARC, American Radiochemicals) were added. Cultures were incubated for 4 h, cells were harvested, and lipids were extracted (33). Aliquots of lipid extracts were analyzed by two-dimensional TLC (HPTLC silica gel 60; Merck) using chloroform:methanol:water (14:6:1, v/v/v) as a mobile phase for the first dimension and chloroform:methanol:glacial acetic acid (13:5:2, v/v/v) as mobile phase for the second dimension (34).

*V. cholerae* strains were inoculated from precultures grown in the same medium. After the addition of 1 μCi of [^14^C] acetate (60 mCi mmol-1) or 0.5 μCi [^14^C] ornithine to each culture, the cultures were incubated overnight. Cells were harvested by centrifugation, washed with 500 μl water once, resuspended in 100 μl water, and then lipids were extracted according to Bligh and Dyer (33). Aliquots of the lipid extracts were spotted on high-performance TLC silica gel 60 plates (Merck, Poole, UK) and separated in two dimensions as described above. To visualize membrane lipids, developed two-dimensional TLC plates were exposed to autoradiography film (Kodak) or a PhosphorImager screen (Amersham Biosciences). The individual lipids were quantified using ImageQuant software (Amersham Biosciences) (8).

The *E. coli* strains expressing the putative OL synthases VC0489 or VCA0646 and an *E. coli* strain harboring an empty vector were grown to an OD of 1.2 at 620 nm in LB medium with kanamycin. Cells were harvested by centrifugation, and lipids were extracted according to Bligh and Dyer (33). Normal phase Liquid chromatography/tandem mass spectrometry analysis (LC-ESI MS) of the lipid extracts was performed using an Agilent 1200 Quaternary LC system coupled to a high-resolution TripleTOF5600 mass spectrometer (Sciex, Framingham, MA). Experimental details can be found in (5).

### Characterization of *V. cholerae* strains

#### Assays for sensitivity to antibiotics

Petri dishes containing 20 ml of LB agar were prepared and overlaid with 5 ml of LB soft agar containing 100 microliters of wildtype cells or mutant strains with an OD of 0.3 at 620 nm. After agar solidification, disk diffusion tests were performed by placing Polymyxin B 300 UI disks onto the soft agar. The inhibition zone’s diameter was determined after growth for 24 h at 30°C. The experiment was repeated three times.

#### Motility Assays

Swim plates (LB medium with 0.3% agar) were point-inoculated with a toothpick and usually incubated for 24 h at 30°C. Photos were taken at timepoint cero, 10 hours and 24 hours. Swimming was assessed qualitatively by examining the circular turbid zone formed by the bacterial cells migrating away from the point of inoculation. The assays were performed with the *V. cholerae* wildtype, the single mutants deficient in ΔVC0489 or ΔVCA0646, and the double mutant deficient in both genes (ΔVC0489, ΔVCA0646).

#### Biofilm formation assays

Biofilm formation was quantified using the method of crystal violet staining, as previously reported, with some modifications (35, 36). Overnight cultures were diluted 1:100 in LB broth with either 0.15 M or 0.0375 M NaCl. The diluted samples (150 μl) were transferred to polyvinyl chloride (PVC) 96-well plates with a “U” bottom (Corning) and incubated under static conditions at 30 °C for approximately 9 hours. The growth of planktonic cells was measured by quantifying the optical density of the static cultures at 600 nm (OD600) using the plate reader Epoch 2. Then, planktonic cells were discarded, and the wells were carefully washed three times with tap water. The biofilms formed in the air-solid-liquid interface were stained with 150 μl of a 0.1% solution of crystal violet dye for 20 minutes. The dye was discarded from the wells, and the wells were washed three times to remove the excess of the dye. The crystal violet stain retained by the biofilms was solubilized in 160 μl of absolute ethanol, and its OD600 was quantified in the plate reader Epoch 2. Biofilm formation arbitrary units were calculated for each well by dividing the OD600 of the crystal violet stain associated with the biofilm with the OD600 of the planktonic cells that grew statically in the same well.

#### *Caenorhabditis elegans* maintenance and survival assays

*Caenorhabditis elegans* WT strain Bristol N2 was maintained at 20°C on MYOB (Modified Youngren’s, Only Bacto-peptone, (37)) agar plates seeded with *E. coli* OP50 (food strain) by standard methods (38). Survival assays were performed as previously described (20). *E. coli* OP50 and *V. cholerae* A1552 wildtype or *V. cholerae* mutant strains were grown overnight in LB broth at 37°C and 30°C, respectively, and then adjusted to an OD600 of 1. One hundred μl of the adjusted culture were spread onto the center of 6 cm MYOB agar plates and incubated for 24 h at the same temperature used for the overnight culture, before adding 30 to 50 synchronized L4-stage worms per plate. Plates were incubated at 20°C and scored for dead worms every day. A worm was considered dead when it no longer responded to touch. The live worms were transferred to a new plate every two days. The missing worms were excluded from the analysis. The survival curves were plotted and analyzed by using the program PRISM, version 9.5.1 (GraphPad Software, San Diego, California USA, www.graphpad.com). Survival curves were considered significantly different from each other when *p* < 0.05.

#### Galleria mellonella survival assays

With bacterial cultures grown to an OD600 of 0.1 in LB medium, serial dilutions in 10 mM MgSO_4_ corresponding to an OD600 of 0.05, 0.025, 0.010, and 0.005 were made. Simple puncture of the larvae, injection of saline solution, or injection of *E. coli* DH5α [17] at an OD600 of 0.1 were used as negative controls. This strain is harmless to *G. mellonella* [18; 19]. Assays were performed by the injection method with *G. mellonella* larvae, and 10 µl of bacterial suspensions were injected into the dorsal region of the third anterior abdominal segment of the larvae with a 31G (gauge) insulin syringe. Each bacterial suspension was analyzed using 15 insect larvae individually placed in Petri dishes without diet and incubated at 30 °C. Mortality was assessed every 24 hours for 5 days after injection. At least three independent experiments were performed.

## ACKNOWLEDGEMENTS

This study was supported by grants from CONAHCyT (237713 and 425886) to CS, and PAPIIT UNAM (IN208116 and IN208319) to CS, and (IA200519 and IA201821) to DZS. We acknowledge technical support by María de los Ángeles Moreno Ocampo and Lourdes Martínez Aguilar.

## Supplementary Figures

**Supplementary Figure 1:** Optimization of the phosphate concentration of the defined growth medium used in this study. (A) Growth of wildtype *V. cholerae* in defined M9 medium with different phosphate concentrations. The symbols are defined in the inset. (B) Two-dimensional TLC separation of [^14^C] acetate-labeled lipids of wildtype *V. cholerae* grown in LB medium. (C) Two-dimensional TLC separation of [^14^C] acetate-labeled lipids of wildtype *V. cholerae* grown in M9 medium supplemented with 64 mM phosphate. (D) wo-dimensional TLC separation of [^14^C] acetate-labeled lipids of wildtype *V. cholerae* grown in M9 medium supplemented with 4.2 mM phosphate. (E) One-dimensional TLC separation of [^14^C] acetate-labeled (a) or [^14^C] ornithine-labeled (b) lipids of wildtype *V. cholerae* grown in M9 medium supplemented with 4.2 mM phosphate. The numbered arrows indicate the first and second dimensions of the chromatography separations. PG: phosphatidylglycerol; PE: phosphatidylethanolamine; CL: cardiolipin; OL: ornithine lipid.

**Supplementary Figure S2. Growth of mutants deficient in the OL synthases VC0489 and VCA0646 is not affected under low salinity conditions**. Single mutants ΔVC0489, ΔVCA0646, and double mutant 2M (ΔVC0489 and ΔVCA0646) were grown in LB medium with varying NaCl concentrations. Standard LB medium contains 10 g NaCl per liter, corresponding to approximately 0.17 M NaCl. (A) 0.15 M NaCl, (B) 0.075 M NaCl, (C) 0.0375 M NaCl. The symbols are defined in the inset of the figure.

